# Exploring protein conformational ensembles using evolutionary conditional diffusion

**DOI:** 10.64898/2026.01.30.702768

**Authors:** Xinyue Cui, Lingyu Ge, Xinguang Yang, Xuhui Li, Dongliang Hou, Xiaogen Zhou, Guijun Zhang

## Abstract

Protein conformational ensembles encode the dynamic landscapes underlying biological function, regulation, and allostery. Accurately reconstructing such ensembles while balancing conformational distributions accuracy and physical plausibility remains a fundamental challenge in structural biology, particularly when dynamic data is scarce. Here, we propose DiffEnsemble, a diffusion-based framework designed for modeling protein conformational ensembles. DiffEnsemble learns latent dynamical representations from static protein structures in the Protein Data Bank, integrated with the structural profile derived from the AlphaFold Protein Structure Database as conditional guidance during the diffusion process. Benchmarking on 72 protein targets from the ATLAS molecular dynamics simulation dataset demonstrates that DiffEnsemble outperforms existing methods, including BioEmu and AlphaFLOW. Compared with AlphaFLOW, DiffEnsemble achieves improvements of 28.9% and 11.3% in Pearson correlation coefficients for ensemble pairwise root mean square deviation and root mean square fluctuation, respectively. Importantly, DiffEnsemble successfully captures the dominant motions for 42% of the targets. These results demonstrate that latent dynamical information embedded in static structural data can effectively support the modeling of protein conformational ensembles.

## INTRODUCTION

Proteins are inherently dynamic molecular machines whose biological functions arise from transitions among interconverting conformational states rather than from a single static structure. These conformational dynamics underpin essential biological processes, including allosteric regulation, enzymatic catalysis, and signal transduction^1^. While computational methods now achieve reliable high-resolution structures for monomeric proteins^2,3^, they still struggle with generating their functional conformational ensembles, which encode the spectrum of dynamic states underlying molecular behavior.

Moving beyond static structural views to address this challenge begins with establishing a well-defined conceptual foundation, which is indispensable for developing any systematic framework that aims to capture conformational diversity. Here, we define a protein conformational “ensemble” as a collection of two or more distinct structural conformations adopted by the same sequence^4^, including functionally relevant alternative and multiple conformations. Alternative conformations usually refer to a limited number of discrete, functionally relevant states of a protein, whereas multiple conformations span a broader range of conformational states. Understanding and modeling such conformational ensembles is central to a fundamental shift in protein science, representing a transition from static depictions to a dynamic view of proteins as molecular machines. Such a focus deepens mechanistic insights into function and establishes a physically realistic framework for linking structure to activity.

The advent of static protein structure prediction methods, exemplified by AlphaFold2^2^ and AlphaFold3^3^, has revolutionized the determination of protein static structures and established the foundation for modeling alternative protein conformations. In recent years, large-scale sampling of multiple sequence alignments (MSAs) based on AlphaFold2 has been regarded as a feasible and effective strategy^4^. Representative methods, including AF-Cluster^5^, AFsample2^6^, and AF2_conformations^7^, sample alternative conformations by clustering MSAs or by employing masking-based and depth-subsampling strategies. Furthermore, the enhancement of evolutionary information within MSAs guided by energetic frustration analysis^8^ and targeted column masking guided by residue flexibility^9^ has been successfully utilized to generate alternative conformations. While these approaches have demonstrated promising performance, they are intrinsically constrained by the availability and quality of MSAs. In particular, for many functionally critical proteins, such as those related to immune-related proteins, coevolutionary information is often weak or unavailable due to the lack of sufficient homologous sequences, resulting in sparse^10^ or noisy MSAs^11^.

Molecular dynamics (MD) simulations^12^, as a classical physics-based approach, can provide multiple conformations at high temporal resolution by explicitly modeling atomic interactions, while their applicability to conformations modeling is limited by high computational cost and strong dependence on the accuracy of the force fields^13^. Recently, generative approaches DiG^14^ and BioEmu^15^ have enabled the efficient modeling of protein multiple conformations by sampling equilibrium protein conformations from implicitly learned Boltzmann distributions^16^, substantially reducing the computational cost of exploring conformational space and alleviating the limitations of MD simulations in terms of accessible timescales and sampling efficiency^13^. These approaches’performance often depends on scarce, high-quality MD simulations for training and validation. AlphaFLOW^17^ further incorporates flow-matching and the AlphaFold2^2^ framework to generate multiple conformations, but it still faces challenges related to computational efficiency and physical plausibility.

Overall, computational modeling of protein conformational ensembles mainly relies on large-scale sampling strategies based on AlphaFold^2,3^, and generative modeling approaches^14,15,17^. Recent assessments from CASP16^4,18^, while acknowledging substantial progress, identify a fundamental and persistent bottleneck in the computational modeling of protein conformational ensembles: the scarcity of high-quality dynamic experimental data^4,19^. This limitation hinders computational methods from generating ensembles with accuracy comparable to experimental benchmarks. Furthermore, it often introduces ambiguity into the evaluation process itself, as the available experimental structures provide only partial coverage of the full conformational landscape^18,20^. When training data are scarce, neural network models struggle to learn the high-fidelity conformational energy landscape, compromising the physical plausibility and reliability of their predictions. Consequently, a promising avenue for progress is to explore how the extensive archive of static protein structures might be strategically utilized to compensate for the current scarcity of dynamic data.

Based on the foregoing analysis, we propose a testable hypothesis that the conformational dynamics of a protein, which are defined by its thermodynamic fluctuations within a single molecular ensemble, can be quantitatively mapped onto the structural diversity accumulated across its homologous family through evolution, particularly the variations observed between different ensembles. This idea builds on the observation that sequence differences among related protein homologs, preserved by natural selection, often help adapt or adjust the same central function, which itself depends on specific shape changes or movements^21,22^. As a result, the different static structures seen across a protein family can be thought of as separate snapshots of the range of shapes the protein needs to adopt to work properly. In short, over evolutionary time, the structural differences among homologs capture the set of possible shapes that a single protein can take on during its normal activity.

In this work, we introduce DiffEnsemble, a diffusion-based framework for modeling protein conformational ensembles, which is grounded in the foregoing hypothesis. The training data for this framework are derived from static structures in the Protein Data Bank (PDB)^23^. To address the limited availability of family-level structural information in the PDB, we further incorporate structural profile features extracted from remote homologs in the AlphaFold Protein Structure Database (AFDB)^24^ as conditional inputs. By integrating structural profile with multiple features such as language model embeddings^25^, the denoising process of the diffusion model is effectively constrained, thereby ensuring the physical plausibility of the generated conformations while enabling a systematic exploration of the protein conformational landscape. We evaluated DiffEnsemble on a fully independent test set consisting of 72 proteins from the ATLAS^26^ MD simulation database, which shares no overlap with the static PDB structures used during training to ensure objectivity and prevent data leakage. Experimental results show that DiffEnsemble substantially outperforms AlphaFLOW^17^, BioEmu^15^, and AlphaFold-based large-scale sampling methods^5-7^ in predicting conformational ensembles. Notably, the framework successfully captures distributional trends of protein motion. This capability not only provides empirical support for our hypothesis that “static structural diversity can be mapped to a dynamic conformational spectrum”, but also highlights the practical utility of DiffEnsemble for studying conformational dynamics.

## MATERIALS AND METHODS

### Data Set

The PDB^23^ compiles three-dimensional structures of proteins resolved under various experimental conditions, encompassing both multiple conformational states of individual proteins and structural diversity among homologous proteins. Their collective distribution captures a set of accessible, functionally relevant conformational states within the protein conformational energy landscape. Inspired by this insight, we expect to leverage the extensive accumulation of static structures in the PDB to compensate for the scarcity of dynamic data and to further uncover the diversity of protein conformational states. In this study, we constructed a dataset capturing the spectrum of structural variations from homologous sequences in the PDB^23^.

Specifically, we collected all protein structures (both monomers and complexes) deposited in the PDB before January 1, 2024, and subsequently split all complexes into their individual monomeric units. To reduce redundancy and group homologous structures, proteins were clustered using MMseqs2^27^ at an 80% sequence identity threshold (detailed in **Supplementary Note S1**), yielding 65,327 initial clusters. The centroid of each cluster was designated as the representative sequence and structure. To further ensure inter-cluster non-redundancy and enhance model generalization, these representatives were refined using a more stringent 30% sequence identity cutoff, resulting in 25,016 distinct structural clusters. For each cluster, a shared structural profile was constructed by retrieving remote homologous sequences from the AFDB^24^. Clusters lacking corresponding structural profiles were excluded, yielding a final dataset comprising 19,399 clusters (see **Supplementary Figure S1** for details). A temporal split was then applied using January 1, 2023, as the cutoff date. Clusters deposited before this date were assigned to the training set, and those released thereafter were used for validation.

To evaluate the ability of DiffEnsemble to extract dynamic properties from static structural databases (PDB), we benchmarked its performance using an MD simulation dataset that is completely independent of the training data. Following the protocol established by AlphaFLOW^17^, our initial test set included 82 protein targets from the ATLAS dataset^26^ released after May 1, 2019. To ensure computational feasibility and accommodate hardware memory constraints, five proteins with long sequences (PDB IDs: 6smsA, 6lrdA, 6l3rE, 6l8sA, and 6oz1A) were excluded. Additionally, five other targets (PDB IDs: 6lusA, 6yhuB, 7buyA, 7jrqA, and 7k7pB) were removed owing to the unavailability of the structural profile features from the AFDB. Consequently, a final curated test set of 72 protein targets (**Supplementary Table S1 and Figure S2**) was employed for all subsequent analyses.

### Overview of DiffEnsemble

DiffEnsemble is a computational framework specifically developed for modeling protein conformational ensembles. Unlike most methods that rely on scarce MD simulation data, DiffEnsemble is built upon the hypothesis that static structural diversity can be mapped onto a dynamic conformational spectrum. It directly infers protein dynamics from static PDB structures and incorporates structural profiles constructed from the AFDB as conditional information. **Figure 1** shows an overview of the DiffEnsemble protocol. Starting from a single input conformation, DiffEnsemble represents the protein as a residue-level graph. Node features integrate residue type, ESM embeddings, and structural profile, while edge features encode geometric information, including C_α_ coordinate frame, backbone orientation, and side-chain torsion angles. These representations are subsequently processed by an SE(3)-equivariant network^28^. Through iterative conditioned denoising, DiffEnsemble progressively transforms the initial conformation into a physically plausible conformational ensemble (For more details, please refer to the “**Model Architecture**” section).

**Figure 1.**
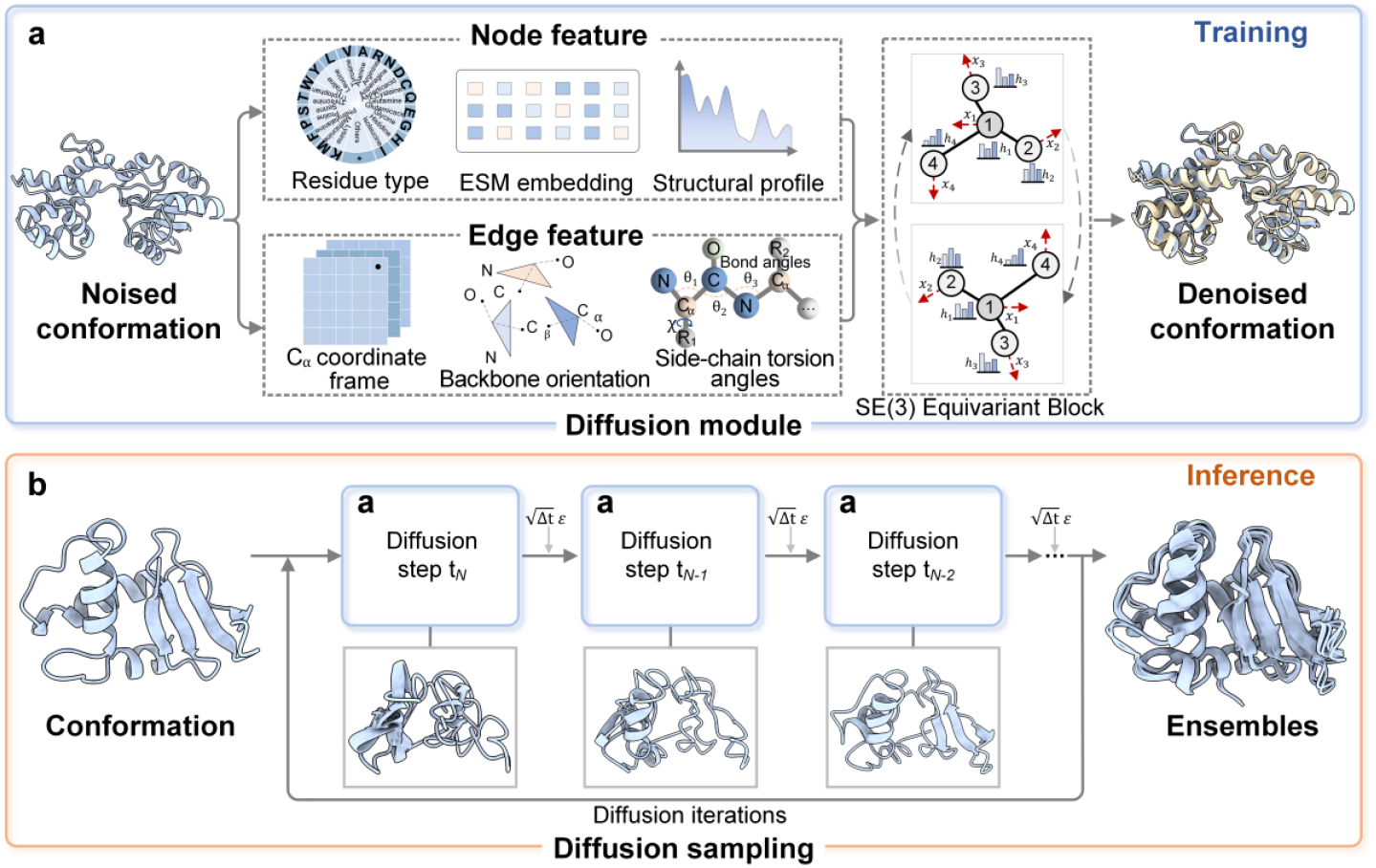
The pipeline of DiffEnsemble includes both training and inference stages. a, Training stage, each protein conformation is initially encoded as a residue-level graph. Residue type, ESM embedding, and structural profile features are incorporated as conditioning information to constrain the denoising process. The geometric states defined by C_α_ coordinate frame, backbone orientations, and side-chain torsion angles. b, Inference stage, the model begins with a conformation and iteratively applies the learned denoising and reverse diffusion steps using the pre-trained network (a), ultimately generating protein conformational ensembles. During each reverse step, a stochastic perturbation is scaled by the square root of the timestep Δt . The noise vector *ε* is sampled from a standard normal distribution.

### Feature Extraction

In this study, proteins are represented by node features that include residue type, ESM embedding, and structural profile, while edge features encode C_α_ coordinate frame, backbone orientation, and side-chain torsion angles.

#### Node Feature

To capture the latent structural and functional information encoded in protein sequences, each residue node is represented by contextual embeddings derived from the ESM-2 language model^25^. These embeddings encode long-range dependencies and evolutionarily conserved patterns within the amino acid sequence, providing rich semantic priors. In DiffEnsemble, the sequence embeddings serve as conditional inputs that guide the diffusion process, enabling the generation of physically plausible conformational ensembles.

To supplement the limited family-level information available in the PDB^23^ and to provide effective constraints that guide the diffusion model toward physically plausible conformations, we constructed structural profile features for each protein. Remote homologous sequences are identified from the UniRef90^29^ database using Jackhmmer^30^, and their corresponding three-dimensional structures are retrieved from the AFDB^24^. After aligning these structures to the target protein, pairwise inter-residue distance distributions are extracted and discretized over a fixed range, yielding a representation that provides a structural prior guiding the diffusion process (**Supplementary Figure S3**).

#### Edge Feature

The edge feature around each residue is described through a combination of C_α_ coordinate frame, backbone orientation, and side-chain torsion angles. Backbone orientation is defined as a residue-centered reference frame that encodes the relative positioning and direction of the backbone atoms (N, C_α_, C) for each residue, providing a rotation- and translation-invariant description of the local backbone geometry. Side-chain torsion angles are parameterized by five rotatable torsion angles ( χ_1_,χ _2_,χ_3_,χ_4_,χ_5_ ) and two symmetry-related angles 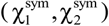. These geometric descriptors enable the model to account for local steric constraints and side-chain orientations, which are critical for maintaining physically realistic conformations during the diffusion process. A detailed description of all features and their dimensions is provided in **Supplementary Table S2**.

### Model Architecture

DiffEnsemble is formulated as a continuous-time diffusion model operating on a residue-level graph neural network with built-in SE(3) equivariance^28^. Information propagation is implemented through tensor products of irreducible representations, following the equivariant framework provided by the e3nn library^31^. Node and edge features are augmented with a sinusoidal embedding of the diffusion time and encoded by separate multilayer perceptrons (MLPs). Residue interactions are restricted to spatial neighbors within a fixed cutoff distance 15Å, with a bounded number of neighbors per residue to balance computational efficiency and memory usage. Node representations are iteratively updated through equivariant tensor product layers, and the final embeddings are used to predict residue-level rigid-body transformations together with side-chain torsion angles.

Model training is performed under a score-based diffusion formulation as a stochastic differential equation (SDE) framework^32^. Perturbed protein conformations are generated by applying morph-like^28^ deformations to native structures, and the learning objective is defined as a conditional denoising score-matching task, in which corrupted input conformations are mapped back toward their corresponding native conformations. For each perturbed conformation, node and edge features are constructed to capture local geometric relationships. Specifically, residue type, ESM embedding, and the structural profile are incorporated as conditioning information for the diffusion model, providing contextual constraints that guide conformational adjustment, while edge features derived from the three-dimensional protein geometry serve as input representations. These features are jointly processed by an SE(3)-equivariant graph neural network^28^.

The loss function is defined as an equally weighted combination of the deviations in protein rotation, translation, and side-chain torsion angles, with the detailed formulation provided in **Supplementary Note S2**. Model parameters are optimized using the Adam^33^ optimizer with parameters *β*_1_ =0.9, *β*_2_ =0.99, *ε* =1e-8. The initial learning rate is set to 0.001 and is dynamically adjusted using a plateau-based scheduler with a patience of 10 epochs. Training is performed with a batch size of 32 on an NVIDIA A100 Tensor Core GPU.

During inference, protein structures predicted by AlphaFold3^3^ are used as the initial noisy conformations. Guided by the conditioning information, the model performs 20 steps of reverse diffusion to refine the conformation progressively. To facilitate broad exploration of the energy landscape and prevent the final conformations from being trapped in a local minimum, a small noise 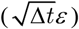 is added to the denoised conformation. By repeating the inference procedure multiple times, the model generates an ensemble of physically plausible protein conformations that capture the intrinsic structural variability of the protein.

### Evaluation Metrics

To rigorously evaluate the quality of the generated conformational ensembles, we assess their performance through the integration of ensemble similarity, distributional accuracy, and physical plausibility. All evaluations are performed with MD simulation ensembles serving as a reference. Definitions of all evaluation metrics are provided in **Supplementary Note S3**.

Ensemble similarity is quantified by Pearson correlation coefficients between the predicted and reference ensembles, calculated from the mean pairwise RMSD^17^ and atom-wise RMSF^34^, in which the mean pairwise RMSD characterizes the conformational diversity of the generated ensembles, whereas RMSF captures atomic positional fluctuations, revealing regions of high flexibility.

Distributional accuracy is assessed by comparing the distributions of C_α_ (glycine for C_β_ ) coordinates projected onto principal component analysis (PCA) space between predicted and reference MD simulation ensembles. The 2-Wasserstein distance (W_2_-dist)^17^ was used to quantify the minimal transport cost between the predicted and reference distributions, providing a quantitative measure of ensemble similarity. In addition, the ability of the model to capture dominant conformational motions is evaluated by computing the percentage of principal component pairs with a cosine similarity greater than 0.5 (%PC-sim > 0.5), which is defined as successful recovery of the principal motions of the ensemble^17^.

Physical plausibility is further evaluated using the radius of gyration^35^ ( R_g_ ), a geometric measure of global protein compactness defined as the root-mean-square distance of all atoms from the center of mass. Larger R_g_ values correspond to more extended conformations, whereas smaller values indicate more compact structures.

## RESULTS AND DISCUSSION

### Performance Evaluation on the ATLAS Benchmark Dataset

To assess the performance of DiffEnsemble in modeling protein conformational ensembles, we benchmarked it against two representative generative model baseline approaches. The comparison included BioEmu^15^ and AlphaFLOW^17^. All methods were evaluated on 72 non-redundant proteins from all-atom MD simulations in the ATLAS^26^ dataset. To ensure statistical fairness, we generated 250 conformations per target for each method. Model performance was assessed across three primary dimensions involving conformational ensemble similarity, distribution accuracy, and structural plausibility. Additional comparisons with AlphaFold-based sampling methods, including AF-Cluster^5^, AFsample2^6^, and AF2_conformations^7^, are provided in **Supplementary Tables S3 and Figures S4-S6**.

As shown in **Figure 2a-c**, DiffEnsemble demonstrates strong performance in reproducing protein conformational ensemble similarity, achieving high Pearson correlations with MD-derived reference ensembles for both pairwise RMSD (*r* = 0.58) and RMSF (*r* = 0.59). To further evaluate whether DiffEnsemble accurately captures the distributional characteristics of protein conformational dynamics, we projected the C_α_ coordinates of each ensemble onto the top two principal components (PC1 and PC2) to evaluate their conformational distributions in a reduced-dimensional space. The differences between predicted and MD-derived reference ensembles were quantified using the 2-Wasserstein distance in PCA space (MD PCA W_2_-dist; **Figure 2d**). Results indicate that the conformational distributions generated by DiffEnsemble are the most similar to the reference distributions compared with baseline methods.

**Figure 2.**
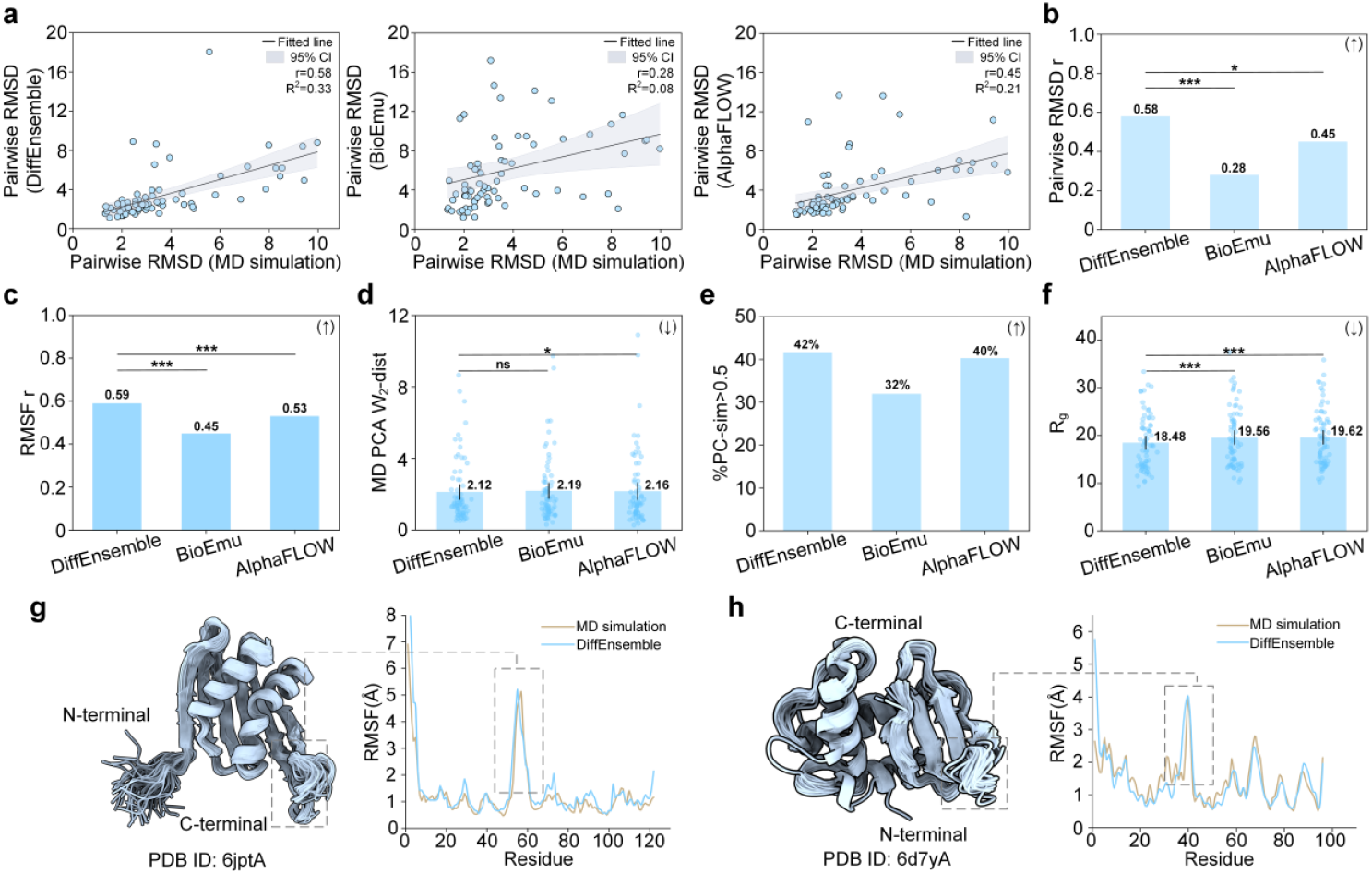
Performance of DiffEnsemble on the ATLAS^26^ benchmark dataset. a, Pearson correlation of pairwise RMSD between predicted and reference ensembles (pairwise RMSD r), with 95% confidence level (CI). b-c, Pearson correlation of pairwise RMSD and RMSF between predicted and reference ensembles (RMSF r), where higher values indicate better agreement (↑). Statistical significance of correlation differences was assessed using Steiger’s Z-test^36^. d, 2-Wasserstein distance (W_2_-dist) between predicted and reference ensemble distributions in PCA space, where lower values indicate better performance (**↓**). e, Percentage of targets whose predicted ensembles achieve a cosine similarity greater than 0.5 to the reference ensembles in the PCA space (%PC-sim>0.5). f, Radius of gyration ( R_g_ ) of predicted ensembles. g-h, Residue-level RMSF analysis of conformational ensembles generated by DiffEnsemble. The bar height indicates the mean; error bars indicate the 95% CI. The MD simulations were treated as the ground truth ensembles and served as the reference for comparison. Statistical significance was determined by first assessing the normality of the paired differences using the Shapiro-Wilk test^37^. For data exhibiting a normal distribution, a two-sided paired t-test^38^ was used. Conversely, non-normally distributed data were evaluated using a two-sided Wilcoxon signed-rank test^39^. Significance levels are denoted as follows: ns, *P* ≥ 0.05; \**P* < 0.05; \*\**P* < 0.01; and \*\*\**P* < 0.001.

In addition, we calculated the proportion of targets exhibiting a cosine similarity greater than 0.5 (%PC-sim > 0.5) between the predicted and reference ensembles distributions (**Figure 2e**). Existing literature^17,40^ reports that a cosine similarity greater than 0.5 indicates successful modeling of dominant collective motions within the conformational ensemble. From this perspective, DiffEnsemble successfully captures the dominant motions for 42% of the targets, highlighting its capability in modeling collective motions underlying protein conformational dynamics. **Figure 2f** further compares the mean radius of gyration^35^ ( R_g_ ) of conformational ensembles predicted by different methods. DiffEnsemble exhibits a significantly lower mean R_g_ than other methods, demonstrating that it generates more compact and physically realistic conformations.

The performance of DiffEnsemble across multiple evaluation metrics highlights its effectiveness in modeling conformational ensembles and provides empirical support for our underlying hypothesis. This improvement can be attributed to the effective integration of static structural datasets with relevant conditional information. The data constructed from the PDB^23^, which aggregates protein structures accumulated over long timescales, enables the model to capture both the evolutionary conservation and the intrinsic dynamic properties of protein structures during training. In addition, the structural profile constructed by searching remote homologous structures from AFDB^24^ offers two key advantages. First, it enables mining the common dynamic characteristics within protein families, which can conditionally guide the reverse diffusion process toward biologically plausible and functionally relevant conformational states. Second, the AFDB encompasses a broader range of conformational states, serving as a valuable complement to the family-level structural information available in the PDB^23^. Leveraging these strengths, DiffEnsemble demonstrates superior generalization capabilities compared with other methods.

Protein dynamics are characterized not only by the timescale of fluctuations (the kinetic component), but also by the amplitude and directionality of these fluctuations (the structural component)^41^. Therefore, to validate the model’s performance, we assessed the dynamical characteristics of the predicted ensembles by analyzing the residue-level RMSF for two representative examples within the test set, using the reference MD simulation ensemble as the benchmark. As illustrated in **Figures 2g** and **2h**, we examined the human PAC3 homodimer^42^ (PDB ID: 6jptA), a key assembly chaperone in the 26S proteasome. Accurate modeling of its conformational dynamics is essential for elucidating intermediate states during proteasome biogenesis. Notably, in the highly flexible region spanning residues 45-60, the ensemble RMSF values predicted by our method closely align with those derived from MD simulations, demonstrating strong consistency in local flexibility estimation.

We further evaluated the model using the CdiA toxin domain from Pseudomonas aeruginosa^43^ (PDB ID: 6d7yA), corresponding to the C-terminal region of the full-length protein. In this case, DiffEnsemble accurately captures large fluctuations in structurally variable regions, reflecting the inherent flexibility of bacterial toxin domains. This result validates the DiffEnsemble’s ability to reproduce residue-level dynamics and provide mechanistic insights into how CDI toxins recognize target cells and how cognate immunity proteins neutralize their activity. While no literature currently validates whether the identified high-probability flexible regions arise from ligand binding or other factors, their high consistency with MD simulation results provides important clues for further investigations into functional states.

### Stereochemical Plausibility of DiffEnsemble-generated Ensembles

We evaluated the stereochemical plausibility of the generated protein conformational ensembles using Ramachandran plot analysis. By examining the statistical distributions of backbone ϕ-ψ dihedral angles, this analysis provides a well-established criterion for assessing whether predicted conformations satisfy fundamental stereochemical properties, thereby offering a stringent test of local backbone quality. Specifically, we compared five representative methods, including BioEmu^15^, AlphaFLOW^17^, AF-Cluster^5^, AFsample2^6^, and AF2_conformations^7^ (**Figure 3** and **Supplementary Figure S7**), for three classes of protein structures: all-α helices, all-β sheets, and mixed α/β folds. To avoid biases associated with atypical backbone geometries, glycine and proline residues were excluded from the analysis. Across all evaluated methods, the predicted ϕ-ψ angle distributions are predominantly concentrated within high probability regions consistent with their corresponding secondary structure elements, indicating that these approaches generally preserve essential backbone stereochemical constraints.

**Figure 3.**
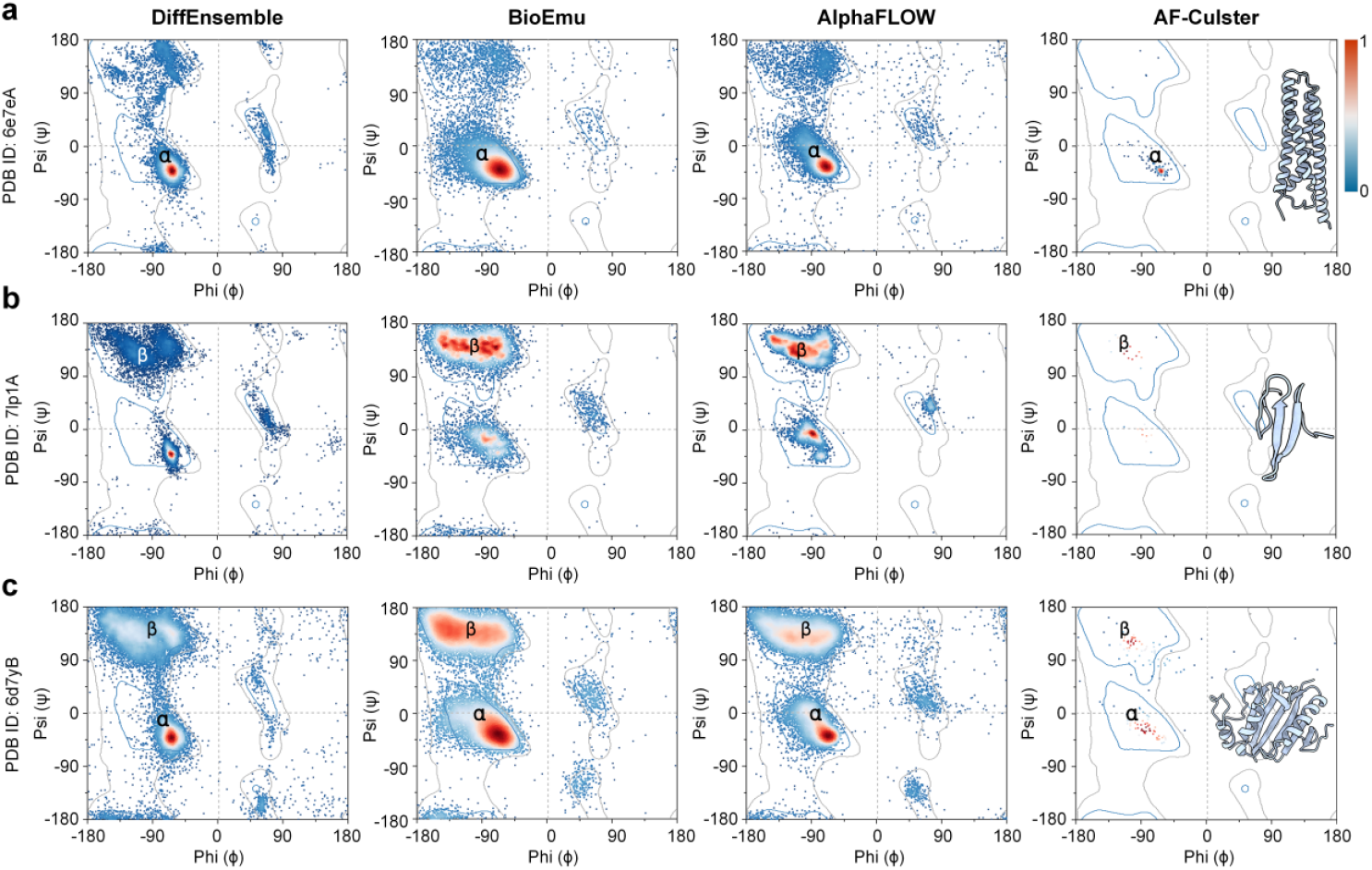
Ramachandran plots illustrating the backbone ϕ-ψ distributions of the conformational ensembles predicted by different methods (250 conformations per target). The color scale ranges from blue (low sampling density) to red (high sampling density). a, Example of all-α protein. b, Example of an all-β sheets protein. c, Example of a mixed α/β folds protein. In the Ramachandran plot, the blue contour-enclosed region represents the favored conformational space (ϕ/ψ dihedral angle combinations), while the gray contour-enclosed area denotes the allowed.

In contrast, clear differences emerge between AlphaFold-based approaches and generative models in terms of Ramachandran plot coverage and distribution patterns. Methods employing modifications to coevolutionary information in multiple sequence alignments to induce conformational diversity (e.g., AF-Cluster^5^) produce ϕ-ψ angle distributions that are markedly more concentrated. This behavior reflects inherent limitations of clustering and input perturbation strategies in capturing broad conformational variability. In contrast, diffusion-based models, including BioEmu^15^, AlphaFLOW^17^, and DiffEnsemble, exhibit substantially broader coverage in Ramachandran space, enabling the sampling of a richer set of local backbone conformational states and demonstrating enhanced conformational exploration capability. It should be noted that diffusion-based generative models, while expanding coverage of conformational space, also sample a small fraction of conformations within sterically disallowed regions of the Ramachandran plot. This observation indicates that extensive exploration of conformational space increases the likelihood of accessing low-probability backbone geometries, particularly in highly flexible or transient structural regions. It also points to a promising direction for future refinement of diffusion-based generative models, in which additional stereochemical or physically motivated constraints may be introduced to reduce sampling in sterically disallowed regions while preserving conformational diversity.

### Effect of Sampling Scale on Conformational Ensemble Performance

The robustness of DiffEnsemble with respect to sampling scale was systematically evaluated by varying the number of predicted conformations across five ensemble sizes (250, 200, 150, 100, 50). The details are provided in **Supplementary Tables S3-S7**. As shown in **Figure 4**, DiffEnsemble exhibits stable performance across all sampling sizes. This low sensitivity to the number of generated conformations demonstrates remarkable robustness in its predictive behavior. Moreover, we observe that for a given method, optimal performance according to different evaluation metrics often occurs at distinct sampling scales. This metric-dependent behavior underscores a broader challenge in the field: the lack of a standardized, biologically grounded framework for the systematic evaluation of protein conformational ensembles.

**Figure 4.**
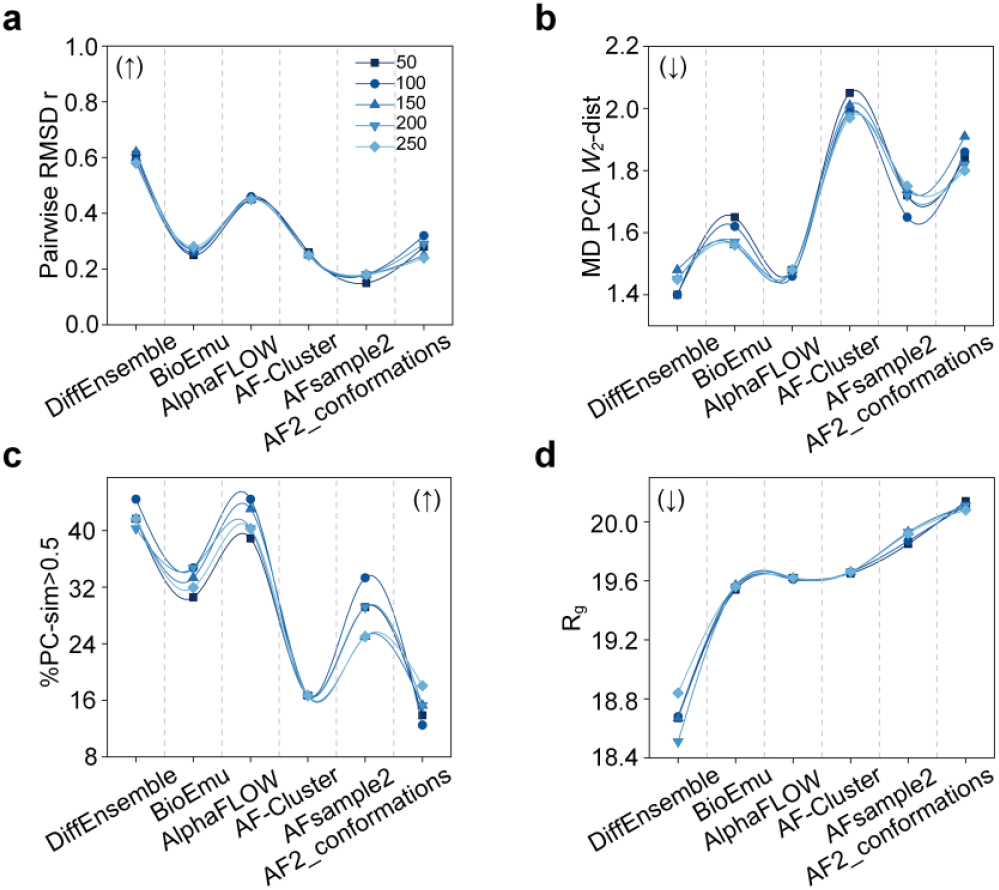
Performance of different methods in generating conformational ensembles across varying numbers of conformations and evaluation metrics, where (↑) denotes that higher values correspond to better performance and (↓) denotes that lower values correspond to better performance. Additional evaluation metrics are provided in **Supplementary Figure S6**.

### Ablation Study

To systematically evaluate the contribution of key conditioning information to the overall performance of DiffEnsemble, we performed validation from two complementary perspectives: structural priors (structural profile) and sequence evolutionary information (ESM embedding). Structural profile constructed from remote homologous proteins captures family-level conformational diversity and provides biologically meaningful conditions during ensembles generation. In parallel, ESM embedding encodes evolutionarily conserved information that is closely associated with functionally relevant conformational dynamics. We performed systematic ablation studies on the ESM embedding and structural profile modules, assessing their impact on model generalization and ensemble diversity.

As shown in **Figure 5a-5c**, removing either the structural profile (w/o structural profile) or the ESM embedding (w/o ESM Embedding) leads to a substantial degradation in performance across all evaluation metrics. Compared with the full DiffEnsemble model, the absence of structural profile results in the most pronounced decline, with Pearson correlation coefficients for pairwise RMSD and RMSF reduced by 10.3% and 15.3%, respectively. In contrast, excluding ESM embedding causes a smaller but still notable reduction (3.4% for pairwise RMSD and 8.5% for RMSF; **Figure 5a**), indicating that structural profile plays a more dominant role in preserving ensemble-level conformational similarity.

**Figure 5.**
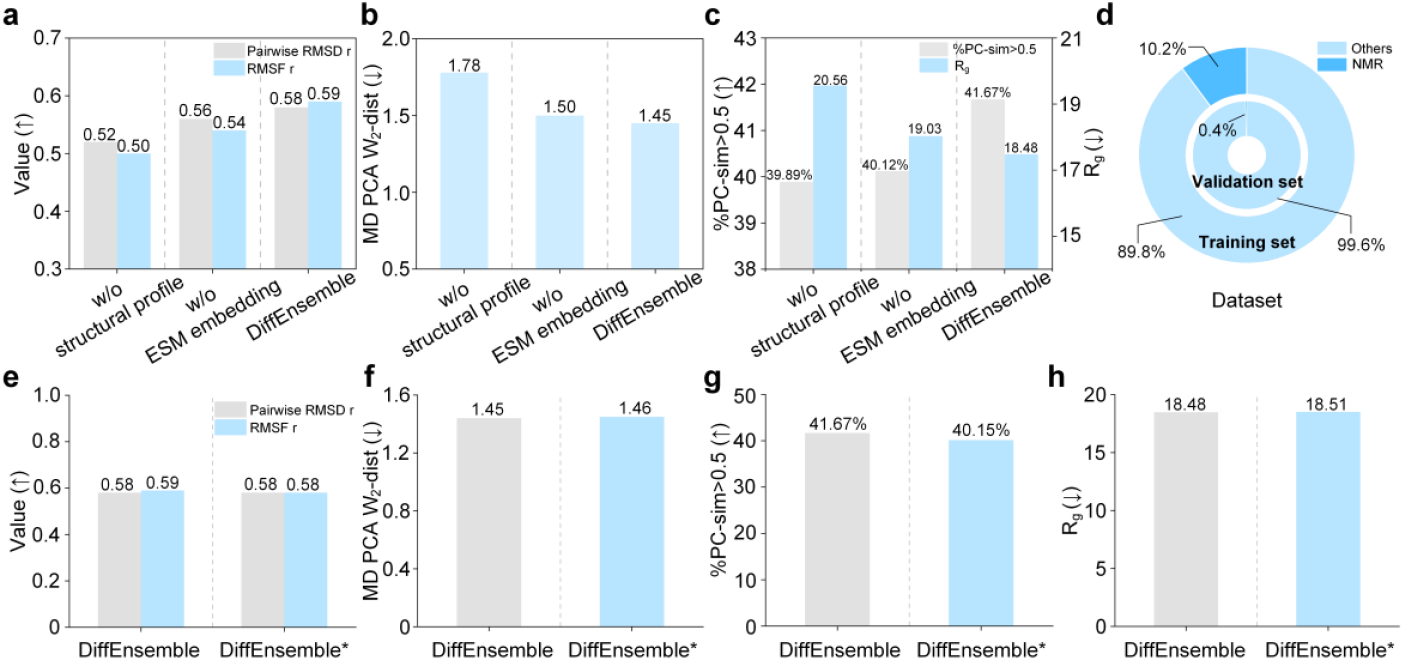
Ablation analysis of DiffEnsemble. a-c, Performance comparison of DiffEnsemble and its variants without using the structural profile (w/o structural profile) and the ESM embedding (w/o ESM embedding). The MD PCA W_2_-dist is reported as the median value. d, Distribution of different data types in the dataset. e-h, Model performance evaluation following the exclusion of Nuclear Magnetic Resonance (NMR) data from both the training and validation sets. DiffEnsemble* denotes the model retrained without using any NMR data.

Consistent with these trends, both ablated variants show reduced distributional accuracy in PCA space, as evidenced by increased 2-Wasserstein distances (MD PCA W_2_-dist; **Figure 5b**) and a decreased fraction of captured dominant motions (%PC-sim > 0.5; **Figure 5c**). In addition, the predicted ensembles become less compact, as evidenced by R_g_ distributions, suggesting that the removal of either structural profile constraints or sequence-derived evolutionary information compromises the physical realism of the generated ensembles. Together, these results demonstrate that structural profile and ESM embedding provide complementary and indispensable conditioning information, jointly enhancing the conformational ensemble similarity, distributional accuracy, and structural plausibility of DiffEnsemble.

To rigorously quantify the contribution of Nuclear Magnetic Resonance (NMR) data^44^ on model performance, we performed a systematic ablation study in which all NMR data were excluded from both the training and validation sets. As detailed in **Figure 5d**, NMR structures account for only 10.2% and 0.4% of the original training and validation sets, respectively. The performance of the resulting model trained without NMR supervision (DiffEnsemble*) is summarized in **Figures 5e-5h**. Compared with the DiffEnsemble model, DiffEnsemble* exhibits a modest decrease across evaluation metrics in the absence of NMR-derived supervision. This performance decrease does not imply that NMR data are unimportant. The limited effect observed is likely due to the relatively small proportion of NMR-derived structures in the training dataset. Importantly, this ablation experiment further demonstrates that DiffEnsemble is capable of effectively extracting intrinsic conformational dynamics from static PDB structures alone, enabling accurate reconstruction of protein conformational ensembles without explicit reliance on experimental dynamical data.

### Performance on Challenging CASP16 Targets

The Critical Assessment of Techniques for Protein Structure Prediction (CASP) provides a rigorous blind benchmark for evaluating the generalization capability of protein modeling methods. In CASP16, a particularly challenging task was introduced involving the inter-domain conformational distributions of Staphylococcal protein A (SpA) in a domain-linker-domain (D-L-D) construct (ZLBT-C)^18^. SpA consists of five domains, two of which are connected by a six-residue linker that is either a wild-type (WT) sequence or an all-glycine (Gly6) variant (**Figure 6**). The WT linker is highly conserved and is thought to introduce an energetic barrier that facilitates binding to host antibodies^18^. Consequently, this protein provides a stringent and biologically meaningful test for assessing a model’s ability to capture inter-domain flexibility and conformational ensembles distributions.

**Figure 6.**
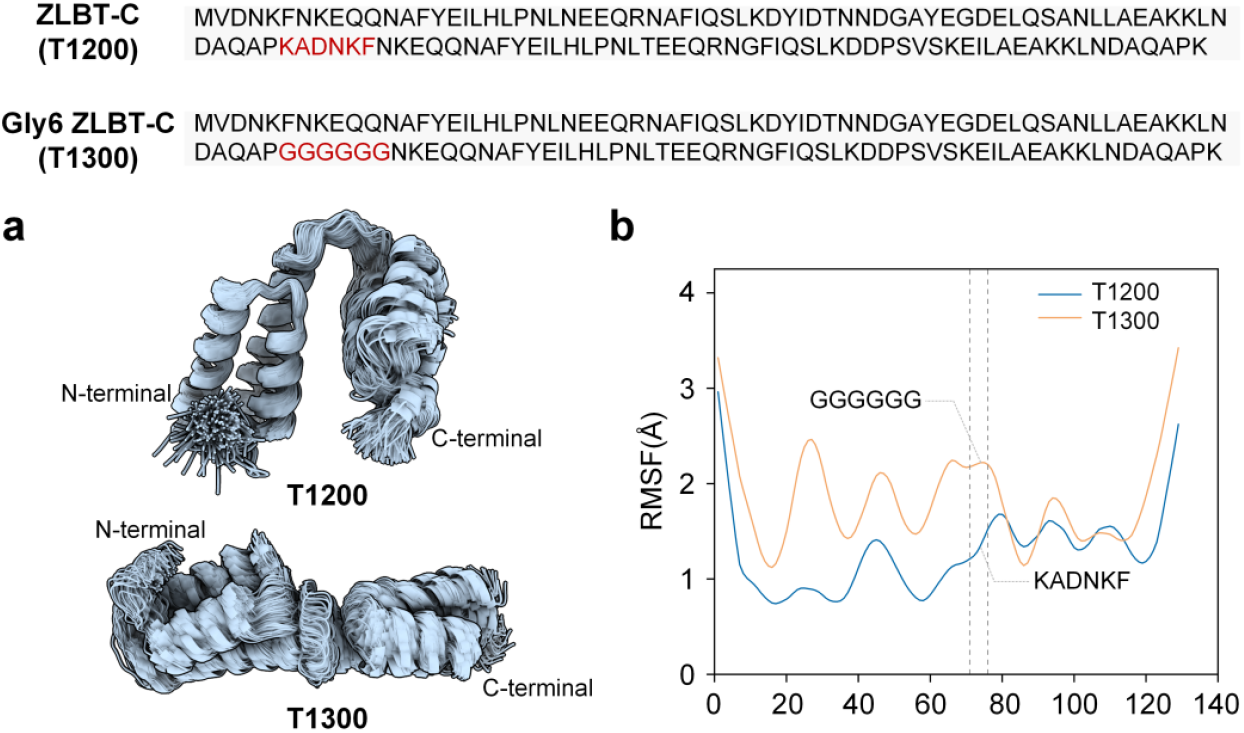
Performance of conformational ensembles prediction on CASP16 targets. a, Protein conformational ensembles for T1200 and T1300 generated by DiffEnsemble. b, Residue-level RMSF of the corresponding conformational ensembles.

To evaluate the performance of DiffEnsemble on these CASP16 ensemble targets, we generated 250 conformations for each target. **Figure 6a** shows that the predicted conformational ensembles for ZLBT-C (T1200) and Gly6 ZLBT-C (T1300) exhibit distinct structural differences. To quantify these dynamic shifts, we calculated the residue level RMSF profiles (**Figure 6b**). The T1300 exhibits a higher mean RMSF (1.83 Å) compared to the T1200 (1.26 Å), indicating that glycine leads to an overall increase in conformational flexibility. Differences in RMSF distributions are also observed in the linker region (residues 71-76), demonstrating a sensitive dynamic response to linker sequence variation. Overall, the T1200 displays more constrained and conservative dynamical behavior, consistent with what is reported in the literature^18^.

Notably, a single metric such as RMSF is insufficient for a comprehensive assessment of ensemble accuracy. In the official CASP16 evaluation, conformational ensembles were assessed using a combination of NMR residual dipolar couplings^45^ (RDCs) and small-angle X-ray scattering (SAXS) profiles^46,47^. However, these metrics were not evaluated in this study because the corresponding evaluation scripts are not publicly available. The final assessment concluded that none were able to recapitulate the conformational ensembles differences between the T1200 and T1300 in the SAXS data^18^. This result highlights the intrinsic difficulty of protein conformational ensembles prediction and underscores the substantial room for improvement in both predictive methodologies and ensembles evaluation metrics. In this context, the successful capture of global dynamic trends by DiffEnsemble demonstrates its robust capacity to characterize the distribution patterns of conformational ensembles.

## CONCLUSIONS

Protein biological function arises from transitions among multiple conformational states. These states occur across a broad range of spatial and temporal scales, and many intermediate states are transient, making them difficult to capture experimentally. Consequently, accurately modeling protein conformational ensembles is a crucial problem and fundamental challenge. Here, we propose DiffEnsemble, a diffusion-based framework for modeling protein conformational ensembles. Unlike most previous generative methods, DiffEnsemble achieves improved accuracy by extracting spatial dynamic information solely from the static PDB^23^ database. To further complement the family-level information absent in the PDB, the model incorporates the structural profile as a condition within the diffusion process, which helps mitigate the generation of stereochemically implausible conformations. Evaluations on an independent MD simulation benchmark^26^ demonstrate that our method outperforms state-of-the-art methods in terms of ensemble similarity, distributional accuracy, and conformational plausibility. These results provide further evidence to support our proposed hypothesis that static structural diversity can be mapped to a dynamic conformational spectrum.

Despite the encouraging results of DiffEnsemble in ensemble modeling, there remains room for improvement in its applicability and accuracy. The current framework primarily focuses on the intrinsic conformational space of proteins and does not fully incorporate the cooperative effects of ligands. Given that many proteins function through an induced-fit mechanism, ligand binding can profoundly reshape the energy landscape and trigger specific conformational transitions. Incorporating ligand priors into the diffusion process will be crucial for achieving precise ensemble modeling. In addition, while our model excels at capturing the statistical distributions of conformational ensembles, it still faces challenges in predicting large-scale conformational rearrangements that span long temporal and spatial scales. A more profound challenge lies in the establishment of an evaluation framework. Currently, metrics for assessing protein conformational ensembles are inconsistent and fragmented, and no consensus has yet been reached across the field. In the future, integrating ligand interactions and physical constraints such as energy-based scoring or force-field restraints, combined with the incorporation of MD simulations^12^ and experimental data (e.g., SAXS^46,47^ and RDC^45^), alongside the development of standardized evaluation metrics, could further improve the predictive performance of DiffEnsemble and extend its applicability to complex molecular systems.

## ASSOCIATED CONTENT

### Data Availability Statement

The authors declare that the data supporting the results and conclusions of this study are available within the paper and its Supplementary Information. All comparative methods were evaluated using their publicly available implementations with default hyperparameters. Specifically, we used BioEmu^15^, AlphaFLOW^17^, AF-Cluster^5^, AFsample2^6^, and AF2_conformations^7^. The corresponding source code is available at its respective GitHub repository. The GitHub repository links for all baseline methods are consolidated in **Supplementary Table S8**.

### Supporting Information

(Note S1) Parameter selection for MMseqs2; (Note S2) Loss function for model training; (Note S3) Evaluation metrics; (Table S1) Detailed information for the 72 test datasets; (Table S2) Feature descriptions and feature dimensions; (Tables S3-S7) Performance comparison of different methods on the 72 ATLAS test datasets; (Table S8) Links to the code implementations of different methods; (Figure S1) Dataset processing; (Figure S2) Three-dimensional structures of the test datasets; (Figure S3) Process for generating structural profile; (Figures S4-S6) Performance of different methods; (Figure S7) Ramachandran plot of AFsample2 and AF2_conformations across different protein types.

## AUTHOR INFORMATION

### Authors

Xinyue Cui - College of Information Engineering, Zhejiang University of Technology, Hangzhou 310023, China

Lingyu Ge - College of Information Engineering, Zhejiang University of Technology, Hangzhou 310023, China

Xinguang Yang - College of Information Engineering, Zhejiang University of Technology, Hangzhou 310023, China

Xuhui Li - College of Information Engineering, Zhejiang University of Technology, Hangzhou 310023, China

Dongliang Hou - College of Information Engineering, Zhejiang University of Technology, Hangzhou 310023, China

### Author Contributions

The manuscript was written through the contributions of all authors. All authors have approved the final version of the manuscript.

### Notes

The authors declare no competing financial interests.

## ACKNOWLEDGMENTS

This work was supported by the National Key R&D Program of China (2022ZD0115103), the National Nature Science Foundation of China (62573386, 62203389), the “Pioneer” and “Leading Goose” R&D Program of Zhejiang (2025C01190), the Zhejiang Province High-level Talent Special Support Program (2023R5248), and Fundamental Research Funds for the Provincial Universities of Zhejiang (RF-C2024006).

## Supplementary Information for

### This file includes

- Supplementary Notes S1 to S3
- Supplementary Tables S1 to S8
- Supplementary Figures S1 to S7
- Reference

## Supplementary Notes

### Note S1.MMseqs2 clustering procedure

To reduce redundancy and group homologous sequences, we employed MMseqs^1^ to cluster all protein sequences from the Protein Data Bank^2^ (PDB). The clustering was performed with the following command:

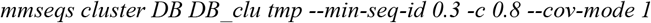

The parameters were set as follows:

- --min-seq-id 0.3: Specifies a minimum sequence identity threshold of 30%, ensuring that only sequences sharing at least 30% identity are clustered together.
- -c 0.8: Sets a minimum alignment coverage of 80%, requiring that the alignment covers at least 80% of the reference sequence.
- --cov-mode 1: Indicates that coverage is calculated for the target sequence, i.e., the aligned portion must span at least 80% of the target.

This configuration maintains a balance between preserving sequence similarity and introducing diversity, which is essential for capturing both conserved and variable structural features.

### Note S2.Loss function for model training

To describe global rigid body motions, the rotation and translation between protein conformations are determined using the Kabsch algorithm^3^. For each residue *i*, let **t**_*i*_ ∈ ℝ ^3^ denote the translational displacement of the noisy protein conformation relative to its native state, representing the translation noise injected during the diffusion process. Correspondingly, 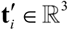 represent the predicted translation vector. For a protein with *N* residues, the translation loss is defined as:

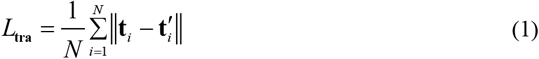

The spatial orientation of residue *i* is parameterized by a rotation matrix **R**_*i*_, with the predicted rotation denoted by 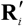 . The rotation loss is defined as follows:

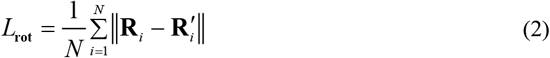

Side-chain torsion angles are defined by five rotatable torsion angles ( χ_1_,χ _2_,χ_3_,χ_4_,χ_5_ ) and two symmetry-related angles 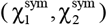. For residue *i* and its corresponding *k* -th ( *m* -th) torsion angle, the loss terms for the five rotatable torsion angles ( *l*_*i,k*_ ) and two symmetry-related angles 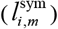 are calculated as follows:

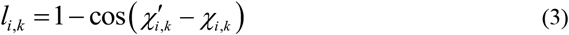

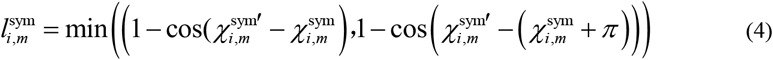

the *χ* ′ represents the predicted torsion angle, while *χ* represents the noise applied during the diffusion process. The total side-chain torsion angles loss ( *L*_tor_ ) is aggregated as:

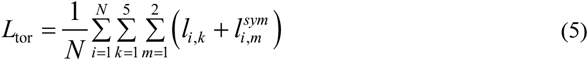

Finally, the training loss for DiffEnsemble is defined as a weighted combination of residue translation, residue rotation, and side-chain torsion angles. For a protein with *N* residues, the total loss is expressed as:

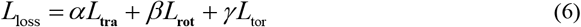

where *α, β, γ* denote the weights of translation, rotation, and torsion angle loss components, respectively (set to *α* = *β* = *γ* = 1 3 in this work).

### Note S3.Evaluation Metrics

We characterized the internal structural diversity by computing the mean pairwise RMSD across all conformer pairs within each ensemble. Subsequently, the Pearson correlation coefficient was calculated between the predicted and MD ensembles based on this metric to assess their alignment in conformational space. The pairwise RMSD^4^ is defined as:

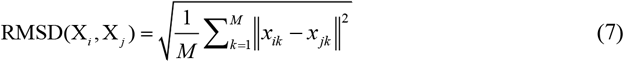

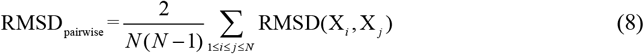

where X_*i*_ and X _*j*_ denote the *i* -th and *j* -th protein conformations within an ensemble of size *N* . Each conformation is represented by the three-dimensional coordinates of corresponding C_α_ atoms, such that *x*_*ik*_, *x*_*jk*_ ∈ ℝ ^*M* ×3^ . Here, *k* ∈{1,…, *M*} indexes the C_α_ atoms, and *x*_*ik*_ denotes the coordinate vector of the *k* -th C_α_ atom in conformation *i* .

The atom-wise RMSF was employed to assess the local conformational variability of each residue. For the *i* -th atom, the RMSF is defined as^5^:

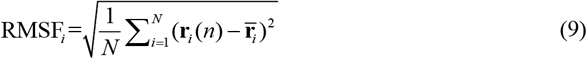

where *N* denotes the total number of conformations in the ensemble,**r**_*i*_ (*n*) represents the position vector of the *i* -th atom in the *n* -th conformation, while **r** denotes the average position vector of the *i* -th atom in the ensemble.

We evaluated the distributional agreement between predicted conformational ensembles and reference MD ensembles by comparing the distributions of C_α_ (glycine for C_β_ ) coordinates in the PCA space. Specifically, the 2-Wasserstein distance (W_2_-dist) was used to quantify the minimal transport cost between the predicted and reference distributions, providing a quantitative measure of ensemble similarity. In addition, the model’s ability to capture dominant conformational motions was assessed by computing the percentage of principal component pairs with a cosine similarity greater than 0.5 (%PC-sim > 0.5), which was defined as successful recovery of the ensemble’s principal motions^4^. Together, these metrics establish a high-resolution benchmark for evaluating the model’s capacity to sample biologically relevant conformational space. The radius of gyration ( R_g_ ) is a geometric measure that quantifies the overall compactness of a protein conformation. It is defined as the root-mean-square distance of all atoms from the center of mass. Larger R_g_ values correspond to more extended conformations, whereas smaller values indicate more compact structures. It is computed as follows^6^:

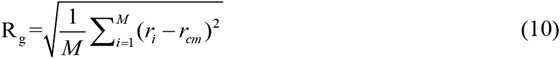

The *M* denotes the total number of atoms in the protein, *r*_*i*_ is the position vector of the *i* -th atom, and *r*_*cm*_ represents the position vector of the center of mass of all atoms.

## Supplementary Tables

**Table S1.** List of 72 ATLAS protein names and lengths. Abbreviations: No., index; Length, the length of the protein sequence.

| No. | Name | Length | No. | Name | Length | No. | Name | Length |
| --- | --- | --- | --- | --- | --- | --- | --- | --- |
| 1 | 5znjA | 567 | 25 | 6ovkR | 219 | 49 | 7aexA | 275 |
| 2 | 6d7yA | 96 | 26 | 6p5hA | 105 | 50 | 7aqxA | 364 |
| 3 | 6d7yB | 155 | 27 | 6p5xB | 457 | 51 | 7asgA | 594 |
| 4 | 6e7eA | 169 | 28 | 6pceB | 70 | 52 | 7bwfB | 92 |
| 5 | 6gusA | 155 | 29 | 6pnvA | 411 | 53 | 7c45A | 302 |
| 6 | 6h49A | 157 | 30 | 6pxzB | 104 | 54 | 7dmnA | 377 |
| 7 | 6h86A | 88 | 31 | 6q10A | 155 | 55 | 7e2sA | 233 |
| 8 | 6iahA | 240 | 32 | 6q9cA | 155 | 56 | 7eadA | 135 |
| 9 | 6in7A | 81 | 33 | 6q9cB | 418 | 57 | 7ec1A | 505 |
| 10 | 6j56A | 129 | 34 | 6qj0A | 412 | 58 | 7jflC | 47 |
| 11 | 6jptA | 122 | 35 | 6ro6A | 58 | 59 | 7la6A | 221 |
| 12 | 6jv8A | 76 | 36 | 6rrvA | 127 | 60 | 7laoA | 271 |
| 13 | 6jwhA | 473 | 37 | 6rwtA | 184 | 61 | 7lp1A | 39 |
| 14 | 6ktyA | 196 | 38 | 6tgkC | 105 | 62 | 7mf4A | 554 |
| 15 | 6l34A | 68 | 39 | 6tlyA | 99 | 63 | 7n0jE | 101 |
| 16 | 6l4lA | 281 | 40 | 6uofA | 119 | 64 | 7nedA | 546 |
| 17 | 6l4pB | 145 | 41 | 6vjgA | 110 | 65 | 7nmqA | 365 |
| 18 | 6ndwB | 178 | 42 | 6xb3H | 241 | 66 | 7onnA | 410 |
| 19 | 6nl2A | 594 | 43 | 6xdsA | 492 | 67 | 7p41D | 448 |
| 20 | 6o2vA | 135 | 44 | 6xrxA | 554 | 68 | 7p46A | 282 |
| 21 | 6o6yA | 440 | 45 | 6y2xA | 235 | 69 | 7qsuA | 374 |
| 22 | 6oddB | 74 | 46 | 6ypiA | 97 | 70 | 7rm7A | 228 |
| 23 | 6okdC | 51 | 47 | 6zslB | 603 | 71 | 7s86A | 99 |
| 24 | 6onoC | 76 | 48 | 7a66B | 86 | 72 | 7wabA | 484 |

**Table S2.** Detailed description of the network’s input features. Where *L* represents the length of the protein sequence.

| Features | Description | Shape |
| --- | --- | --- |
| Residue type | The 20 standard amino acids are encoded in proteins | $L \times 20$ |
| ESM embedding | ESM embedding is a learned representation of protein sequence derived from a deep protein language model. The embedding encodes rich biological information, including evolutionary patterns, structural constraints, and functional characteristics | $L \times 1280$ |
| Structural profile | The structural profile was constructed by statistically analyzing evolutionarily conserved structural features across homologous protein structures from the AlphaFold Protein Structure Database (AFDB) | $L \times L \times 36$ |
| $C_\alpha$ coordinate frame | $C_\alpha$ Coordinates describe the 3D spatial positions (x, y, z) of α-carbon atoms in a protein structure | $L \times 3$ |
| Backbone orientation | Backbone orientation refers to the spatial arrangement of a protein’s main chain (N-$C_\alpha$-C), primarily determined by the dihedral angles $\phi$ (C-N-$C_\alpha$-C) and $\psi$ (N-$C_\alpha$-C-N) | $L \times 2$ |
| Side-chain torsion angles | Side-chain torsion angles ($\chi$ angles) describe the rotational conformations of amino acid side chains in proteins, defining their spatial arrangement relative to the backbone | $L \times 7 \times 2$ |

**Table S3.** Results of 250 conformations predicted by different methods on 72 ATLAS benchmark test sets.

| Methods | RMSD r | RMSF r | MD PCA W<sub>2</sub>-dist |  | %PC-sim>0.5 | R<sub>g</sub> |
| --- | --- | --- | --- | --- | --- | --- |
|  |  |  | Average | Median |  |  |
| DiffEnsemble | 0.58 | 0.59 | 2.12 | 1.45 | 41.67 | 18.48 |
| BioEmu | 0.28 | 0.45 | 2.19 | 1.56 | 31.94 | 19.56 |
| AlphaFLOW | 0.45 | 0.53 | 2.16 | 1.48 | 40.28 | 19.62 |
| AF-Cluster | 0.25 | 0.27 | 2.50 | 1.97 | 16.67 | 19.66 |
| AFsample2 | 0.18 | 0.24 | 2.65 | 1.75 | 25.00 | 19.92 |
| AF2_conformations | 0.24 | 0.27 | 2.74 | 1.80 | 18.06 | 20.08 |

**Table S4.** Results of 200 conformations predicted by different methods on 72 ATLAS benchmark test sets.

| Methods | RMSD r | RMSF r | MD PCA W<sub>2</sub>-dist<br>(Median) | %PC-sim>0.5 | R<sub>g</sub> |
| --- | --- | --- | --- | --- | --- |
| DiffEnsemble | 0.58 | 0.59 | 1.45 | 40.28 | 18.51 |
| BioEmu | 0.27 | 0.45 | 1.57 | 34.72 | 19.56 |
| AlphaFLOW | 0.45 | 0.53 | 1.48 | 40.28 | 19.62 |
| AF-Cluster | 0.25 | 0.26 | 1.98 | 16.67 | 19.66 |
| AFsample2 | 0.18 | 0.25 | 1.72 | 29.17 | 19.93 |
| AF2_conformations | 0.29 | 0.29 | 1.82 | 15.28 | 20.11 |

**Table S5.** Results of 150 conformations predicted by different methods on 72 ATLAS benchmark test sets.

| Methods | RMSD r | RMSF r | MD PCA W<sub>2</sub>-dist<br>(Median) | %PC-sim>0.5 | R<sub>g</sub> |
| --- | --- | --- | --- | --- | --- |
| DiffEnsemble | 0.62 | 0.62 | 1.48 | 41.67 | 18.67 |
| BioEmu | 0.27 | 0.45 | 1.56 | 33.33 | 19.57 |
| AlphaFLOW | 0.45 | 0.53 | 1.48 | 43.06 | 19.62 |
| AF-Cluster | 0.25 | 0.27 | 2.01 | 16.67 | 19.66 |
| AFsample2 | 0.18 | 0.24 | 1.74 | 25.00 | 19.93 |
| AF2_conformations | 0.25 | 0.25 | 1.91 | 15.28 | 20.10 |

**Table S6.** Results of 100 conformations predicted by different methods on 72 ATLAS benchmark test sets.

| Methods | RMSD r | RMSF r | MD PCA W<sub>2</sub>-dist<br>(Median) | %PC-sim>0.5 | R<sub>g</sub> |
| --- | --- | --- | --- | --- | --- |
| DiffEnsemble | 0.59 | 0.60 | 1.40 | 44.44 | 18.68 |
| BioEmu | 0.26 | 0.45 | 1.62 | 34.72 | 19.56 |
| AlphaFLOW | 0.46 | 0.54 | 1.46 | 44.44 | 19.61 |
| AF-Cluster | 0.25 | 0.27 | 1.99 | 16.67 | 19.66 |
| AFsample2 | 0.18 | 0.24 | 1.65 | 33.33 | 19.87 |
| AF2_conformations | 0.32 | 0.33 | 1.86 | 12.50 | 20.11 |

**Table S7.** Results of 50 conformations predicted by different methods on 72 ATLAS benchmark test sets.

| Methods | RMSD r | RMSF r | MD PCA W<sub>2</sub>-dist<br>(Median) | %PC-sim>0.5 | R<sub>g</sub> |
| --- | --- | --- | --- | --- | --- |
| DiffEnsemble | 0.61 | 0.61 | 1.40 | 41.67 | 18.67 |
| BioEmu | 0.25 | 0.45 | 1.65 | 30.56 | 19.54 |
| AlphaFLOW | 0.45 | 0.53 | 1.48 | 38.89 | 19.62 |
| AF-Cluster | 0.26 | 0.27 | 2.05 | 16.67 | 19.65 |
| AFsample2 | 0.15 | 0.24 | 1.72 | 29.17 | 19.85 |
| AF2_conformations | 0.28 | 0.31 | 1.84 | 13.89 | 20.14 |

**Table S8.**
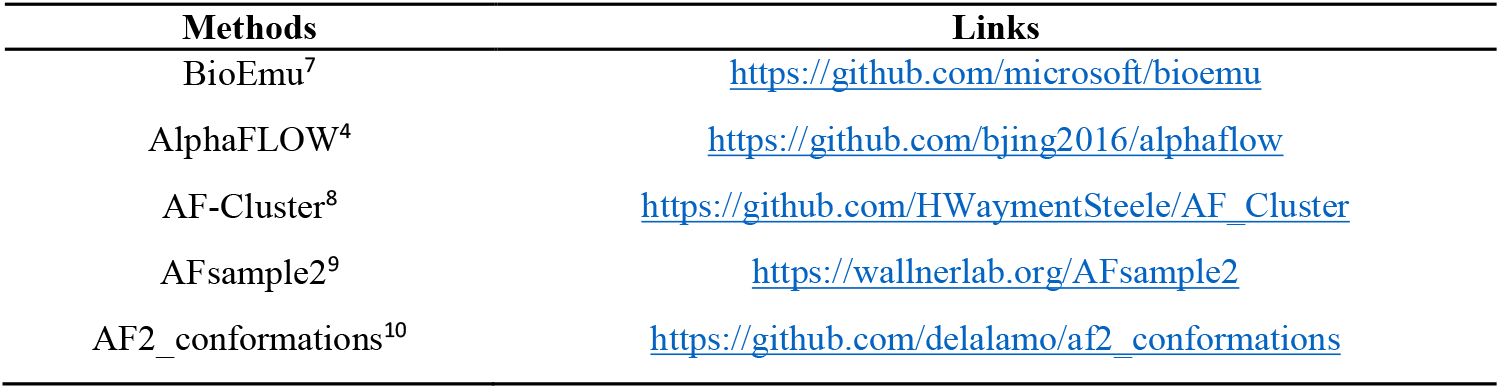
The GitHub repository links for all comparative methods.

| Methods | Links |
| --- | --- |
| BioEmu<sup>7</sup> | https://github.com/microsoft/bioemu |
| AlphaFLOW<sup>4</sup> | https://github.com/bjing2016/alphaflow |
| AF-Cluster<sup>8</sup> | https://github.com/HWaymentSteele/AF_Cluster |
| AFsample2<sup>9</sup> | https://wallnerlab.org/AFsample2 |
| AF2_conformations<sup>10</sup> | https://github.com/delalamo/af2_conformations |

## Supplementary Figures

**Figure S1.**
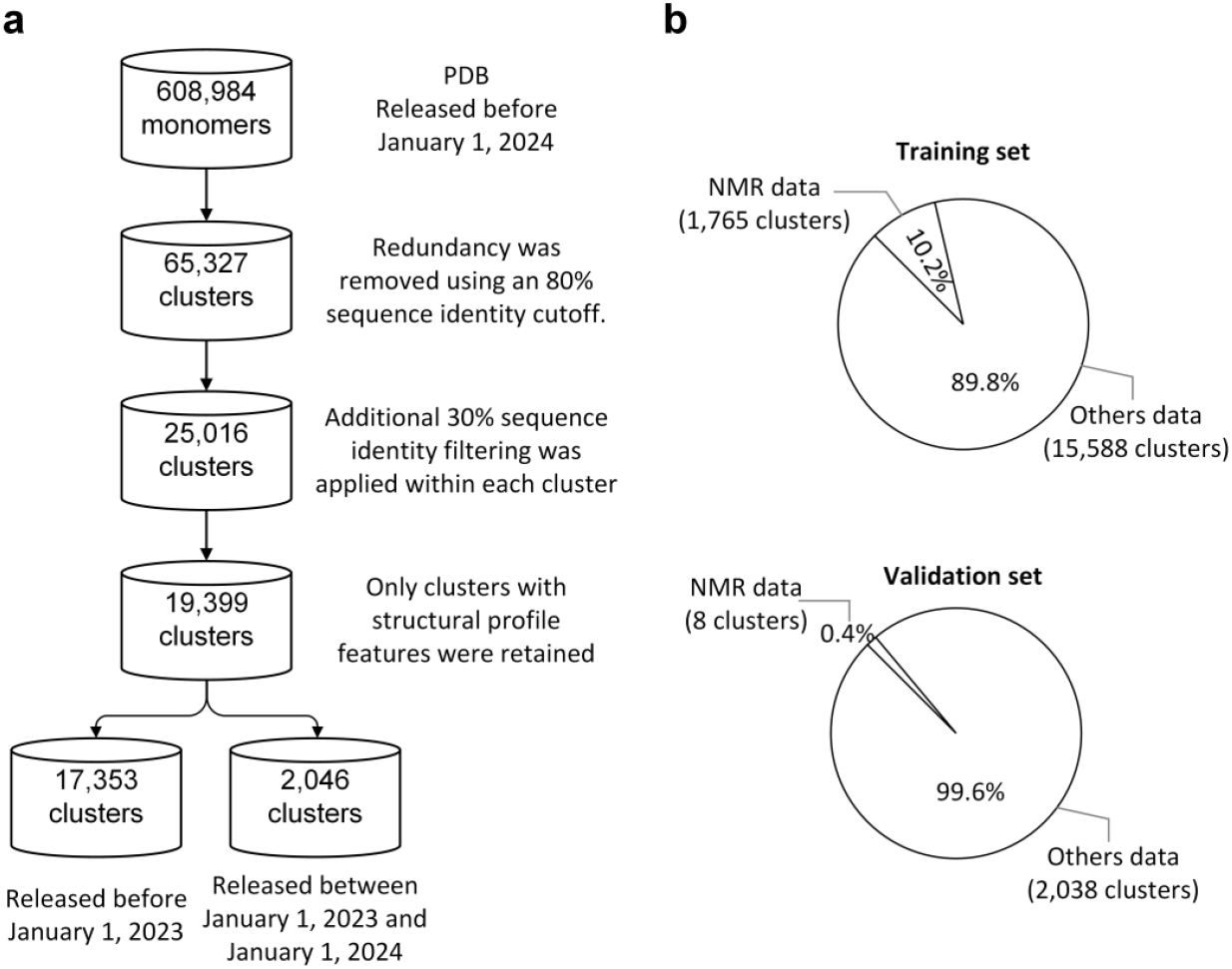
Dataset construction. We collected all protein monomers and complex structures (split into monomers) available in the PDB database as of January 1, 2024, resulting in a total of 608,984 monomers. These structures were clustered at 80% sequence identity, yielding 65,327 clusters. To reduce redundancy, inter-cluster filtering was performed at 30% sequence identity, resulting in 25,016 non-redundant clusters. Clusters with available structural profiles were then selected, resulting in a final dataset of 19,399 structure clusters. Using January 1, 2023, as a temporal cutoff, 17,353 clusters were assigned to the training set and 2,046 to the validation set. b, Distribution of Nuclear Magnetic Resonance (NMR) structures across the training and validation sets.

**Figure S2.**
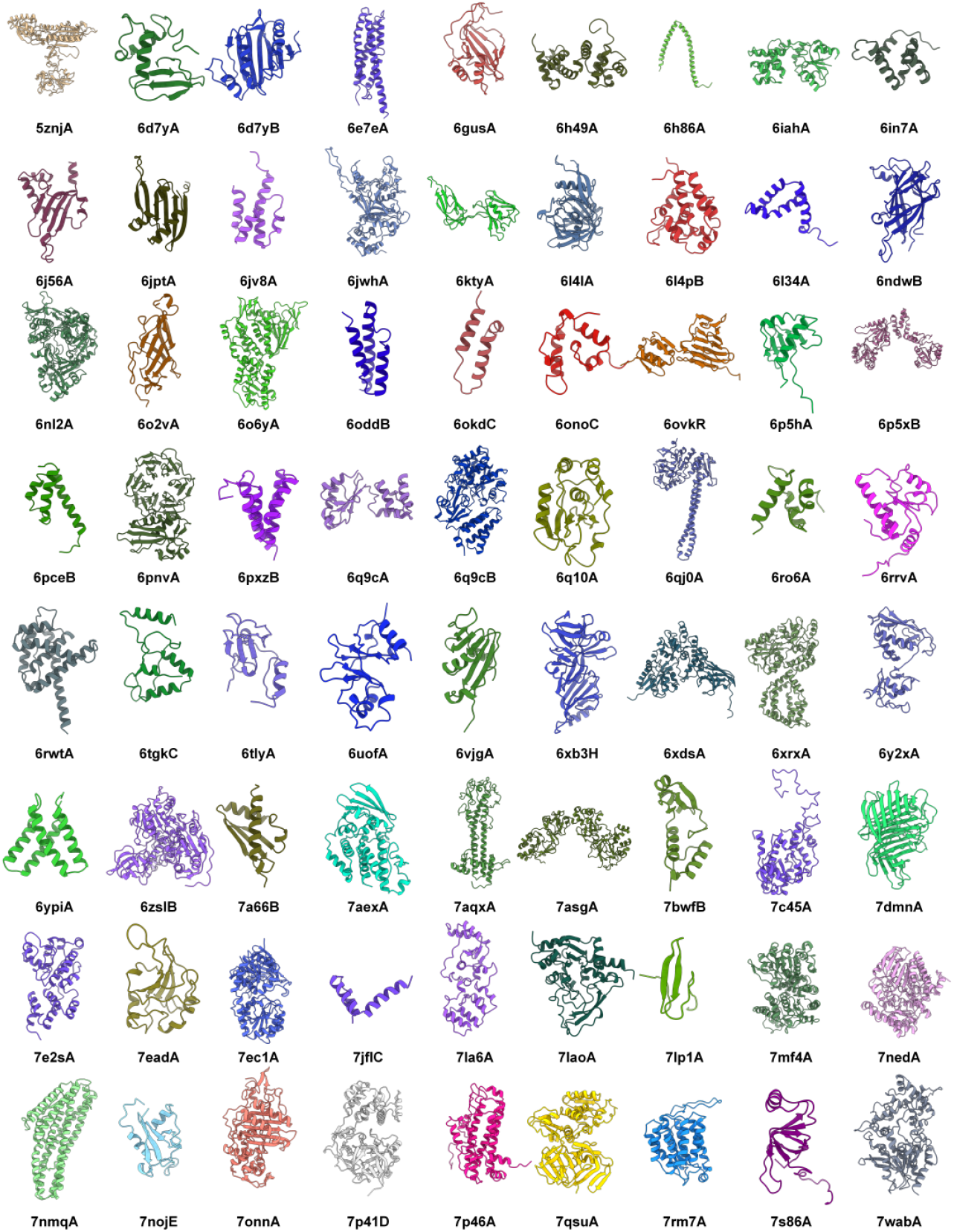
Three-dimensional native conformations of the 72 protein targets.

**Figure S3.**
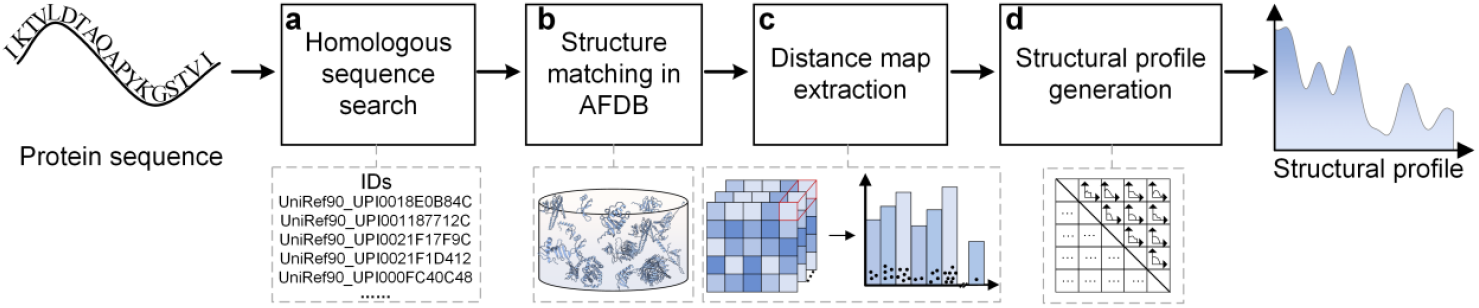
Workflow for generating protein structural profile features. a, Homologous sequence search module, Representative homologous sequences are retrieved from the UniRef90 database using Jackhmmer (E-value < 1e-3). b, High-confidence structural models corresponding to the searched IDs were retrieved from the AlphaFold Protein Structure Database (AFDB). c, Homologous structures were spatially aligned to the target protein. For aligned residues, distance maps between corresponding residues across structures were extracted and discretized into 36 bins ranging from 2.0 Å to 20.0 Å with 0.5 Å intervals. d, The resulting residue-level distributions are aggregated to form the final structural profile features, capturing the evolutionary-informed structural variability of the target protein.

**Figure S4.**
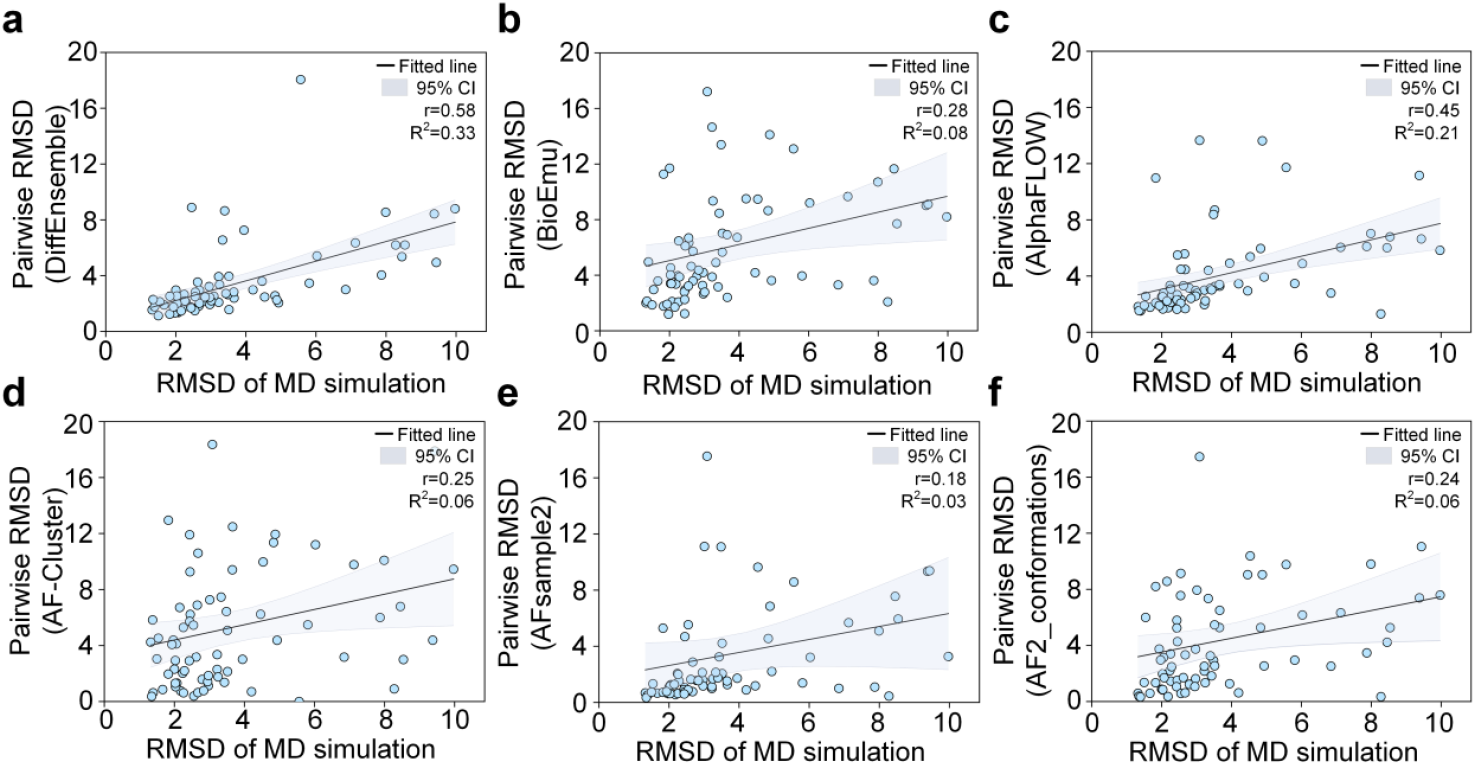
Performance of DiffEnsemble on the ATLAS benchmark dataset. Pearson correlation of pairwise RMSD between predicted and reference ensembles (pairwise RMSD r), with 95% confidence level (CI).

**Figure S5.**
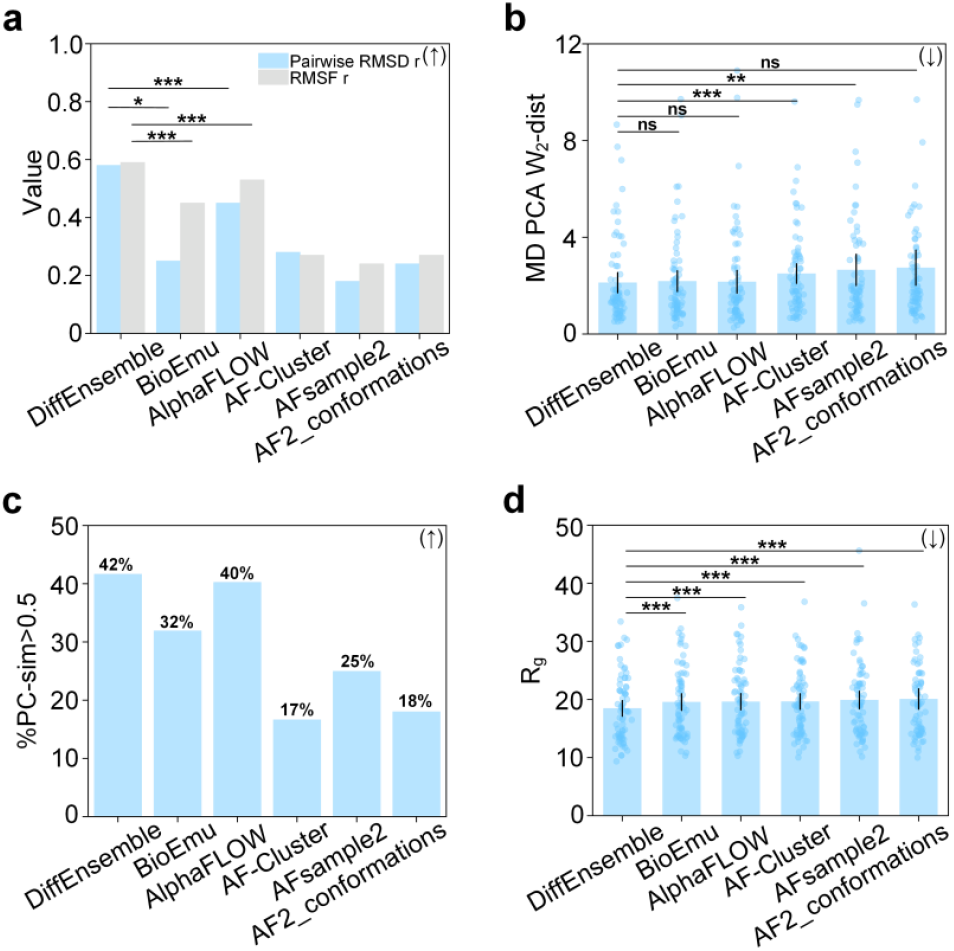
Comparison of different methods on the ATLAS benchmark dataset. a, Pearson correlation of pairwise RMSD and RMSF between predicted and reference ensembles (RMSF r), where higher values indicate better agreement (↑). Statistical significance of correlation differences was assessed using Steiger’s Z-test^11^. b, 2-Wasserstein distance (W_2_-dist) between predicted and reference ensemble distributions in PCA space, where lower values indicate better performance (**↓**). c, Percentage of targets whose predicted ensembles achieve a cosine similarity greater than 0.5 to the reference ensembles in the PCA space (%PC-sim>0.5). d, Radius of gyration ( R_g_ ) of predicted ensembles.

**Figure S6.**
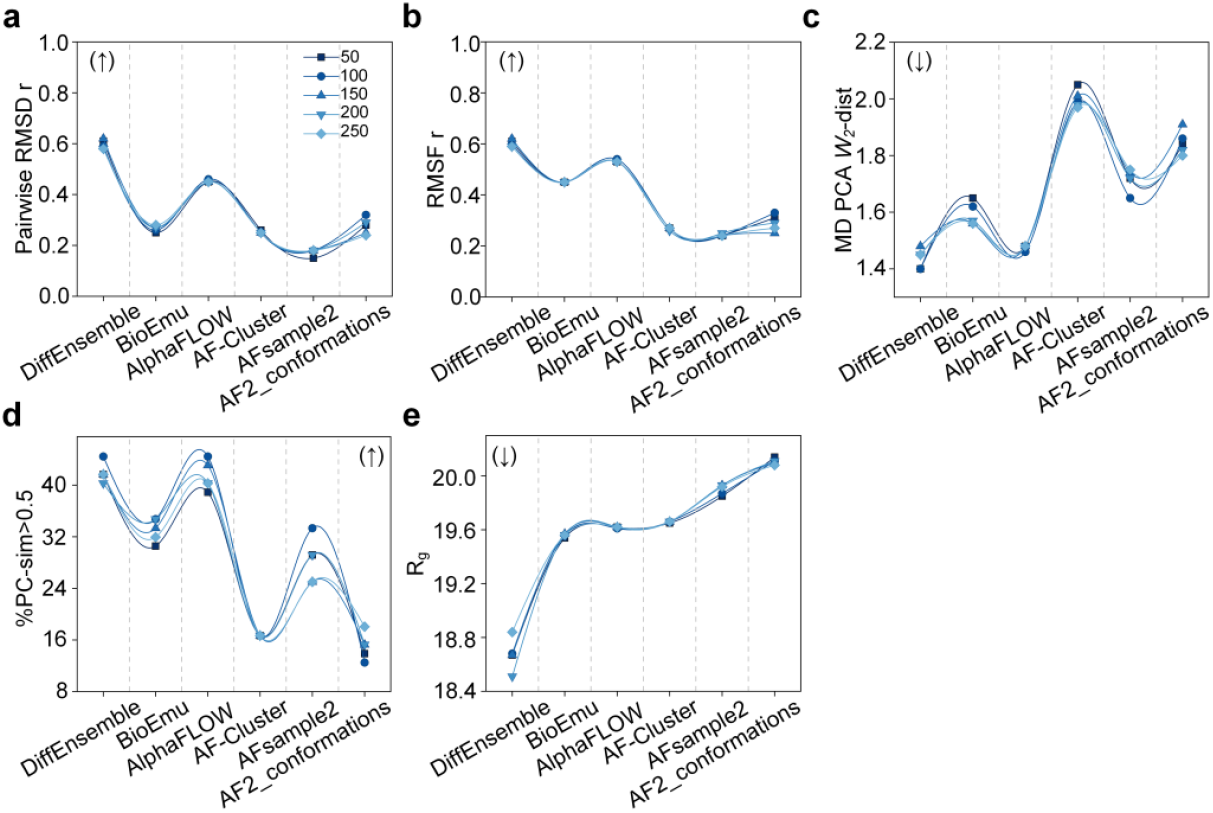
Comparison of model performance across different numbers of generated conformations.

**Figure S7.**
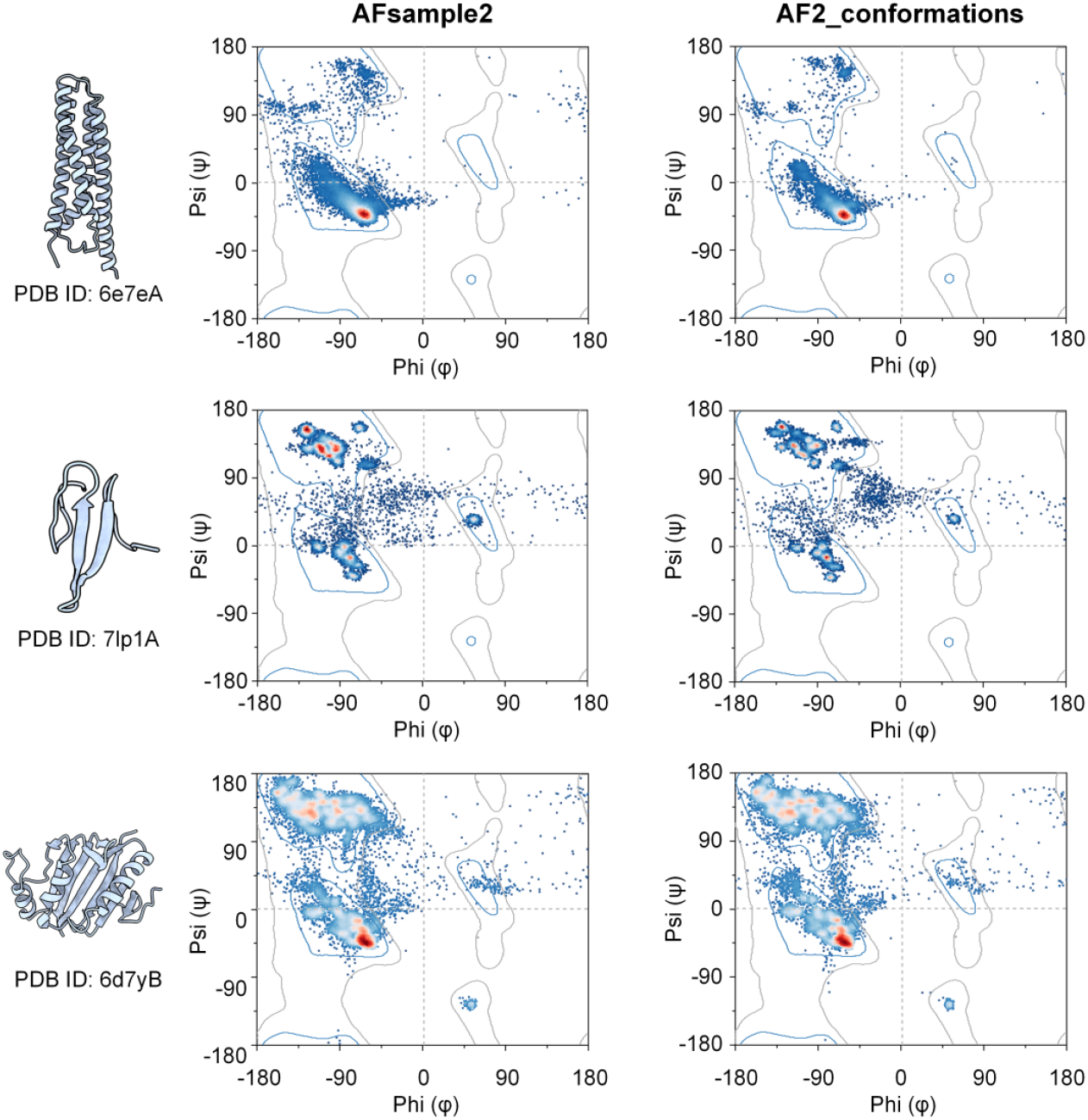
Ramachandran plots of conformational ensembles predicted by AFsample2^9^ and AF2_conformations^10^ for different protein classes.

